# Influenza A virus defective viral genomes are inefficiently packaged into virions relative to wild-type genomic RNAs

**DOI:** 10.1101/2021.05.13.444068

**Authors:** Fadi G. Alnaji, William K. Reiser, Aartjan te Velthuis, Christopher B. Brooke

**Affiliations:** Department of Microbiology, University of Illinois at Urbana-Champaign; Lewis Thomas Laboratory, Department of Molecular Biology, Princeton University; Carl R. Woese Institute for Genomic Biology, University of Illinois at Urbana-Champaign

## Abstract

Deletion-containing viral genomes (DelVGs) are commonly produced during influenza A virus infection and have been implicated in influencing clinical infection outcomes. Despite their ubiquity, the specific molecular mechanisms that govern DelVG formation and their packaging into defective interfering particles (DIPs) remain poorly understood. Here, we utilized next-generation sequencing to analyze DelVGs that form *de novo* early during infection, prior to packaging. Analysis of these early DelVGs revealed that deletion formation occurs in clearly defined hotspots and is significantly associated with both direct sequence repeats and enrichment of adenosine and uridine bases. By comparing intracellular DelVGs with those packaged into extracellular virions, we discovered that DelVGs face a significant bottleneck during genome packaging relative to wild type genomic RNAs. Surprisingly, packaged DelVGs exhibited no signs of enrichment for specific deletion characteristics suggesting that all DelVGs are equally limited in packaging efficiency. Our data provide the first unbiased, high-resolution portrait of the diversity of DelVGs that are generated by the IAV replication machinery and shed light on the mechanisms that underly DelVG formation.

**Importance:** Defective interfering particles (DIPs) are commonly produced by RNA viruses and have been implicated in modulating clinical infection outcomes, hence, there is increasing interest in the potential of DIPs as antiviral therapeutics. For influenza viruses, DIPs are formed by the packaging of genomic RNAs harboring internal deletions. Despite decades of study, the mechanisms that drive the formation of these deletion-containing viral genomes (DelVGs) remain elusive. Here, we used a specialized sequencing pipeline to characterize the first wave of DelVGs that form during influenza virus infection. This dataset provides an unbiased profile of the deletion-forming preferences of the influenza virus replicase. Additionally, by comparing the early intracellular DelVGs with those that get packaged into extracellular virions, we described a significant segment-specific bottleneck that limits DelVG packaging relative to wild type viral RNAs. Altogether, these findings reveal factors that govern the production of both DelVGs and DIPs during influenza virus infection.

## INTRODUCTION

Influenza A virus (IAV) populations are highly heterogeneous and largely consist of virions that lack functional copies of one or more gene segments (1, 2). A major contributor to this heterogeneity is the common presence of defective interfering particles (DIPs) within viral populations. DIPs are virions that harbor a large deletion in one or more genome segments, resulting in an inability to express the full set of viral proteins required for productive replication. DIPs have been demonstrated in numerous contexts to interfere with wild-type virus replication (hence the name), ostensibly either by outcompeting wild type genomes for replication and packaging and/or by triggering innate immune activation (3–7). Recent studies have correlated the abundance of DIPs within clinical samples with severity of both IAV and RSV infection, suggesting a role for DIPs in modulating viral pathogenicity (8, 9). Despite being discovered over 60 years ago, the specific molecular processes that drive DIP formation, as well as the effects of DIPs on the collective behavior and pathogenicity of viral populations remain mysterious (10, 11).

The deletion-containing genomic RNAs carried by DIPs are commonly referred to as defective viral genomes (DVGs) (3). This terminology is complicated for influenza viruses, as a variety of distinct defective viral genome species have been described, including hyper-mutated segments (12) and mini viral RNAs (mvRNAs), which carry enormous deletions and do not get packaged into virions (13). In addition, it is not yet clear that the production of some deletion-containing segments is actually detrimental to viral population fitness (11). Thus, to minimize confusion, we use the term DelVG (Deletion-containing Viral Genome) here to refer to any viral gene segments that contain deletions greater than 10 nucleotides, yet retain the classical viral packaging signals.

Next generation sequencing (NGS) represents a powerful new tool for revealing the specific processes and molecular determinants that underly DelVG formation (14–17). The analysis of large numbers of individual DelVGs can reveal specific patterns that yield mechanistic insight into the formation process (18). IAV DelVGs are typically studied in the context of extracellular DIPs. A potential limitation of this approach for investigating the DelVG formation process is that the requirements of intracellular trafficking and packaging may specifically select for a subset of the total repertoire of DelVGs produced within the cell (19). As a result, DelVGs isolated from extracellular DIPs may not be representative of the full range of DelVG products produced within the infected cell. Such a discrepancy between the DelVGs present within an infected cell and those that get packaged into DIPs was recently described for chikungunya virus (20).

To gain a more accurate, comprehensive understanding of DelVG and DIP formation, we specifically examined the DelVGs that formed *de novo* during the first hours of IAV infection. Careful analysis of hundreds of intracellular and extracellular DelVGs revealed the enrichment of specific sequence elements at deletion breakpoints. We also observed that DelVGs represent a much larger fraction of viral RNAs within the cell compared with what gets packaged, suggesting that IAV DelVGs are packaged much less efficiently than wild type genomic RNAs. While the magnitude of this effect varied somewhat between genome segments, we observed no significant differences in the sizes or locations of packaged DelVG deletions compared with intracellular DelVGs, indicating that the bottleneck on DelVG packaging is not sequence-specific.

## RESULTS

### Intracellular DelVGs generated early during infection are primarily derived from the polymerase and HA segments

Previous examinations of influenza DelVGs have looked specifically at the RNAs that get successfully packaged into virions (DIPs). It is not actually clear how well the packaged DelVGs observed within extracellular DIPs represent the total population of DelVGs produced by the IAV replication machinery. To better understand the full array of deletions commonly generated during IAV infection, and by extension the DelVG formation process, we examined the distributions of deletion breakpoints found within intracellular viral RNAs isolated early during infection.

We infected MDCK-SIAT1 cells at a multiplicity of infection (MOI) of 10 (based on Tissue Culture Infectious Dose (TCID50)) with a recombinant stock of A/Puerto Rico/8/1934 (PR8) grown under conditions that minimize DIP content. We then harvested total cellular RNA at 3 and 6 hours post infection (hpi) in order to capture the intracellular DelVGs produced early during infection. We also extracted viral RNA from supernatants collected at 24 hpi to capture DelVGs that were successfully packaged into DIPs. These RNAs, along with genomic RNA extracted from the infecting viral stock, were used as templates for whole genome RT-PCR and next generation sequencing as we have previously described (17).

To focus our analysis on DelVGs that formed *de novo* during the experiment, we first defined the specific DelVGs present within the inoculum and excluded them from subsequent analyses **(Fig 1A)**. We also only considered deletion junctions if they were represented by >5 reads within a given sample, a cutoff threshold that maximized junction detection albeit with reduced correlation between technical replicates **(Fig 1B)**. We chose to prioritize junction detection sensitivity in this study due to the low copy number and read coverage of *de novo* generated DelVGs early during infection.

**Fig 1.**
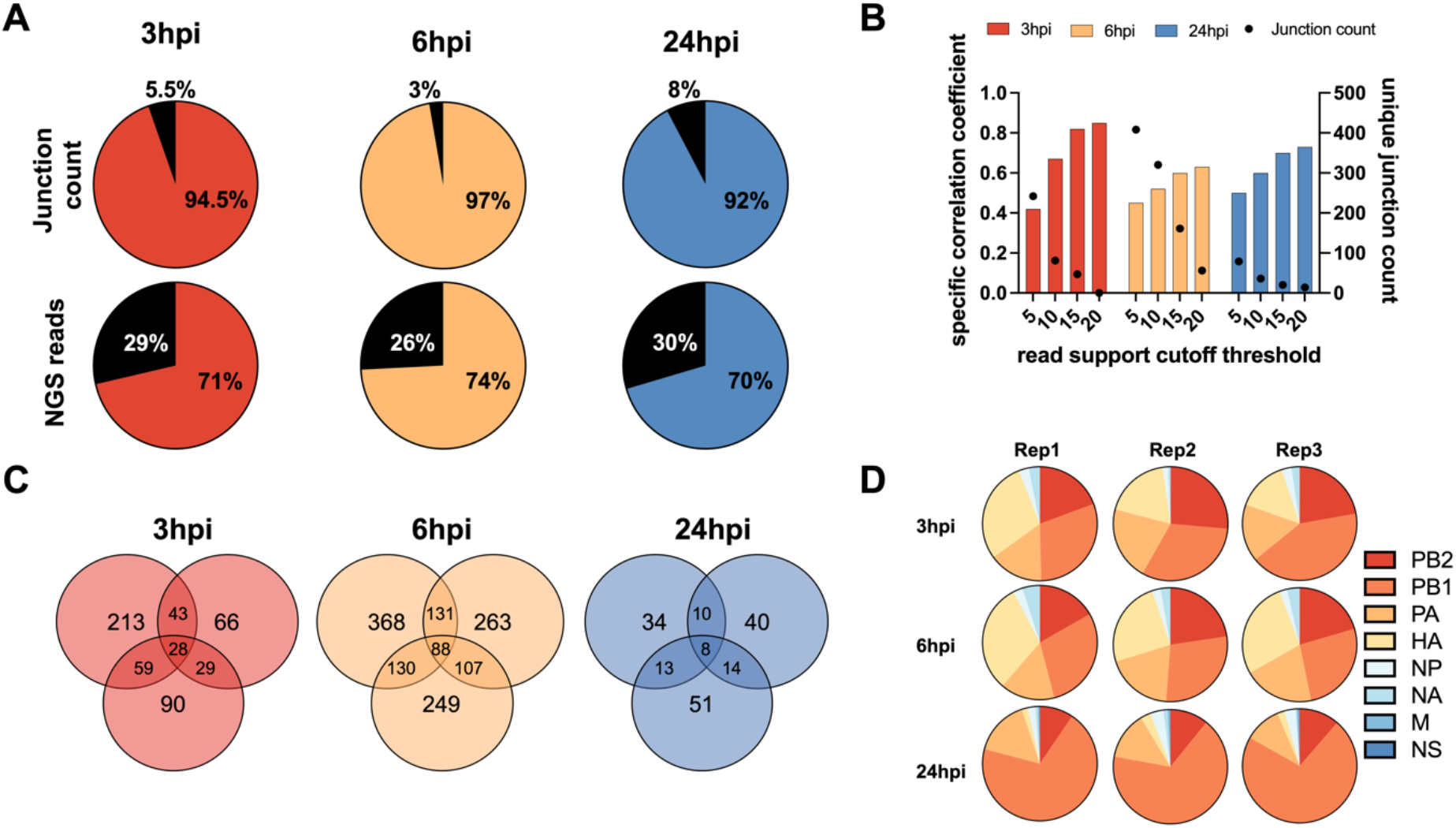
DelVG formation is partially stochastic and biased towards the polymerase and HA segments. **(A)** Pie charts show the percentages of the total normalized junction counts of all segments between the inoculum and viral populations from all time points at the levels of unique junction site (upper panel) and NGS reads (lower panel). The black fractions represent the junctions that were detected in the inoculum. **(B)** Correlation of distinct junctions in the PB2 segment between two technical replicate samples (generated from same RNA sample) using different NGS read cutoff values. **(C)** Venn diagram showing the overlap in specific junctions (for the PB2 segment) between three replicate samples collected at each time point. **(D)** Pie charts show the proportional abundance of normalized junction counts from each genome segment across replicates.

Using this approach, we identified hundreds of distinct DelVGs across all segments except M and NS, in both extracellular and intracellular samples. For each time point, we observed only partial overlap in the specific junctions shared between the three replicates, suggesting significant stochasticity in the specific locations at which deletions form **(Fig 1C)**. When we examined the proportional distribution of junctions across the eight genome segments, we found that these proportions were consistent across replicates, indicating significant and reproducible variation in the intrinsic potential of the individual genome segments to form DelVGs **(Fig 1D)**. Numerous previous studies have shown that the majority of extracellular DelVGs found within DIPs are derived from the three polymerase segments, PB2, PB1, and PA (17, 21, 22). Interestingly, we observed that HA-derived DelVGs were roughly as numerous as polymerase-derived DelVGs at early timepoints within infected cells but only constituted a small fraction of extracellular DelVGs at 24 hpi, suggesting that population of DelVGs packaged into virions may not accurately reflect the relative abundances of DelVGs produced within infected cells.

### DelVG formation is not simply a function of segment length

It is not clear why some segments are more prone to DelVG formation than others, though it has been postulated that this is simply a function of the segment length (23). To directly test this hypothesis, we compared the normalized junction count between DelVGs detected at 6 hpi and a perfect model that assumes a positive correlation between junction count and the proportional length of each segment (**Fig 2A**). While the PB2 and PA segments matched up well with the model predictions, the other segments deviated significantly. These data demonstrate that rates of DelVG formation are not proportional to segment length.

**Fig 2.**
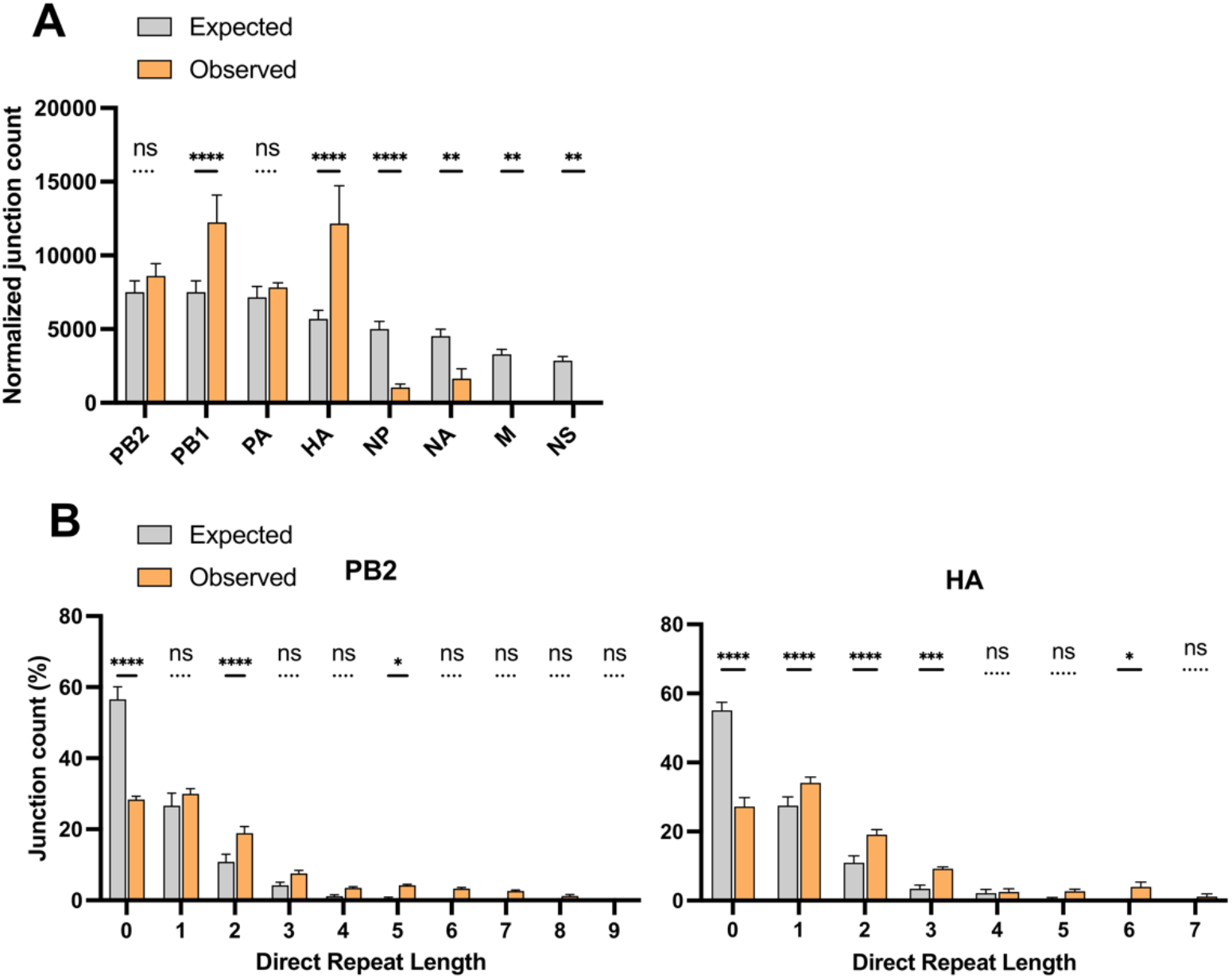
Roles of segment length and direct repeat sequences in DelVG deletion formation. **(A)** Observed normalized junction counts per segment were compared with junction counts predicted by a model (expected) that assumes a simple positive correlation between segment length and DelVG junction count (see materials and methods). **(B)** The numbers of DelVG junctions detected at 6 hpi with no sequence repetition flanking the deletion breakpoints (Direct repeat length = 0), or with repeated sequences of varying length (Direct repeat length = 1-9) for the indicated segments (observed). Expected values plot the numbers of junctions with the indicated repeat lengths predicted from model simulations in which junction formation is random. Junction counts are plotted as a percentage of the total number of DelVGs detected for a given segment. Data are presented as means (n = 3 cell culture wells) ± standard deviations. * P<0.05, ** P<0.01, ***, P < 0.001; ****, P < 0.0001, ns = not significant (Two-way ANOVA).

### Direct sequence repeats and A/U bases are enriched at DelVG deletion junctions

By examining newly produced DelVGs isolated early during infection, we generated an unbiased view of DelVG deletion formation by the IAV replicase. Previous reports have described an enrichment of repeated sequences flanking DelVG deletions (termed “direct repeats”) and have hypothesized these direct repeats may promote DelVG formation by facilitating RNA-dependent RNA polymerase (RdRp) re-engagement during the replication process (15, 24). In support of these previous studies, we observed that deletions lacking direct repeat sequences were significantly less abundant, and a subset of direct repeat sequence lengths were more abundant in early intracellular DelVGs than would be predicted if deletions were completely random **(Fig 2B)**.

We also asked whether specific sequence motifs were enriched at DelVG deletion junctions. Using the 6 hpi samples, we extracted the sequences flanking each deletion breakpoint in the pre-deletion, wild type sequence and calculated the proportion of each nucleotide at each position **(Fig 3)**. To determine whether the observed nucleotide frequencies deviated from what would be expected if deletion formation was sequence independent, we performed the same analysis on three replicate sets of deletions randomly generated *in silico*. Comparison of the observed nucleotide frequencies with the null model predictions revealed clear enrichment of adenosines or uridines at the 5’ deletion breakpoint position (labeled “J” in figure 3). We also detected enrichment of either adenosines or uridines at the position immediately upstream of both the 5’ and 3’ breakpoints (position 4 in both R1 and R3). Finally, we observed enrichment of either cytidines or guanosines within the R2 region downstream of the 5’ breakpoint. Altogether, these results suggest that DelVG formation is facilitated in part by the presence of direct sequence repeats and nucleotide composition surrounding the junction sites.

**Fig 3.**
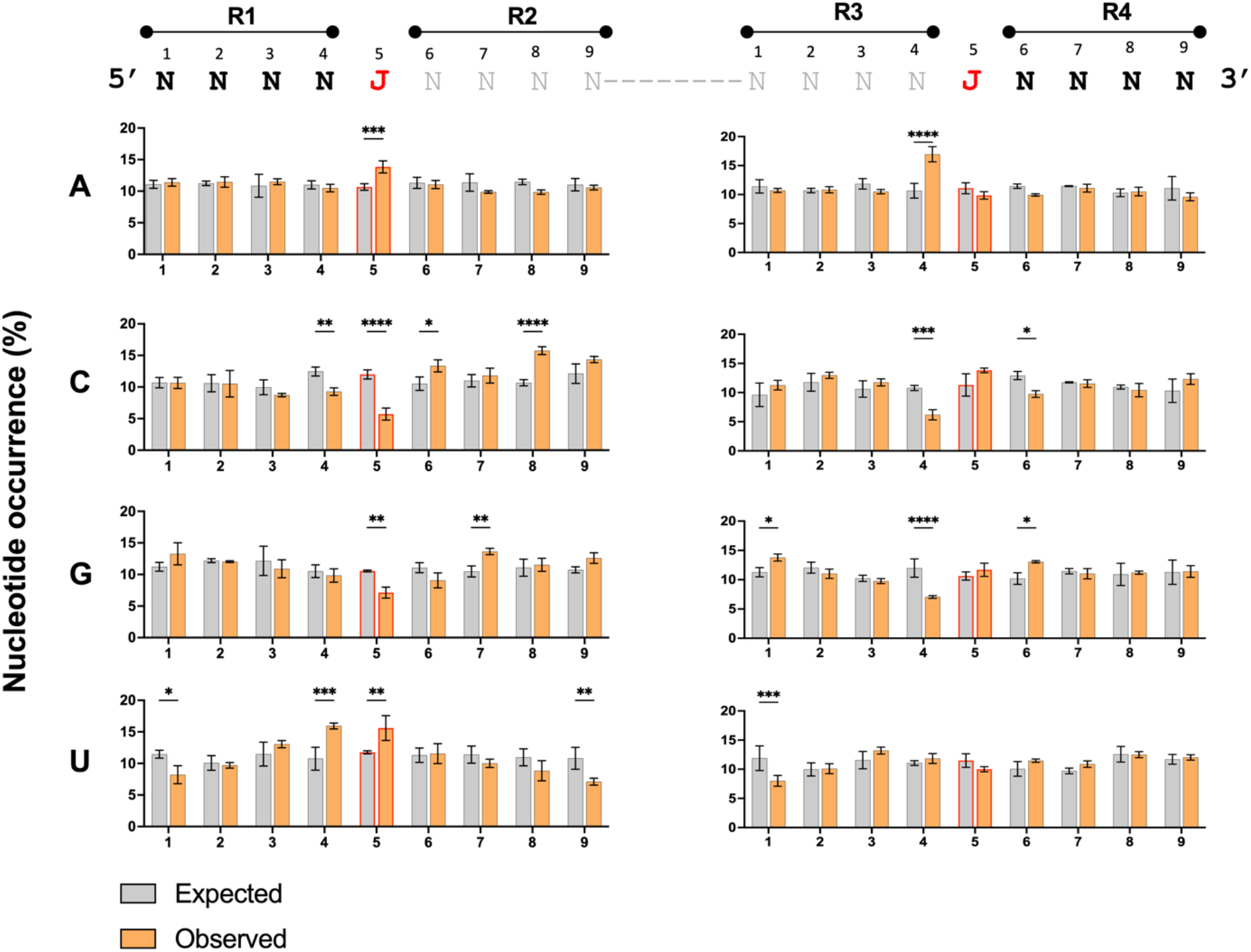
Enrichment of specific nucleotides at positions flanking DelVG deletions. The 4 nucleotides flanking DelVG junctions were numbered and divided into four regions (R1-R4) and the percentage occurrence of each nucleotide was calculated at each site within each region. The junction nucleotides immediately flanking the deletion are indicated by the red “J”‘s at position 5 on the left and right. The grey nucleotides flank the junctions in the progenitor sequence but are lost through deletion in the actual DelVG sequence. The percentage of nucleotide occurrence (observed in PB2 6 hpi) at each site was plotted against a random control (expected). Data are presented as means (n = 3 cell culture wells) ± standard deviations. * P<0.05, ** P<0.01 ***, P < 0.001; ****, P < 0.0001 (Two-way ANOVA), ns = not significant.

### Intracellular DelVGs are inefficiently packaged

Our finding that HA-derived DelVGs make up a large fraction of intracellular but not extracellular DelVGs suggested that some DelVGs might be packaged into virions more efficiently than others. To further investigate this, we compared both the normalized numbers of unique junctions and read support for DelVGs from each segment between intracellular (3 and 6 hpi) and extracellular (24 hpi) populations **(Fig 4A)**.

**Fig 4.**
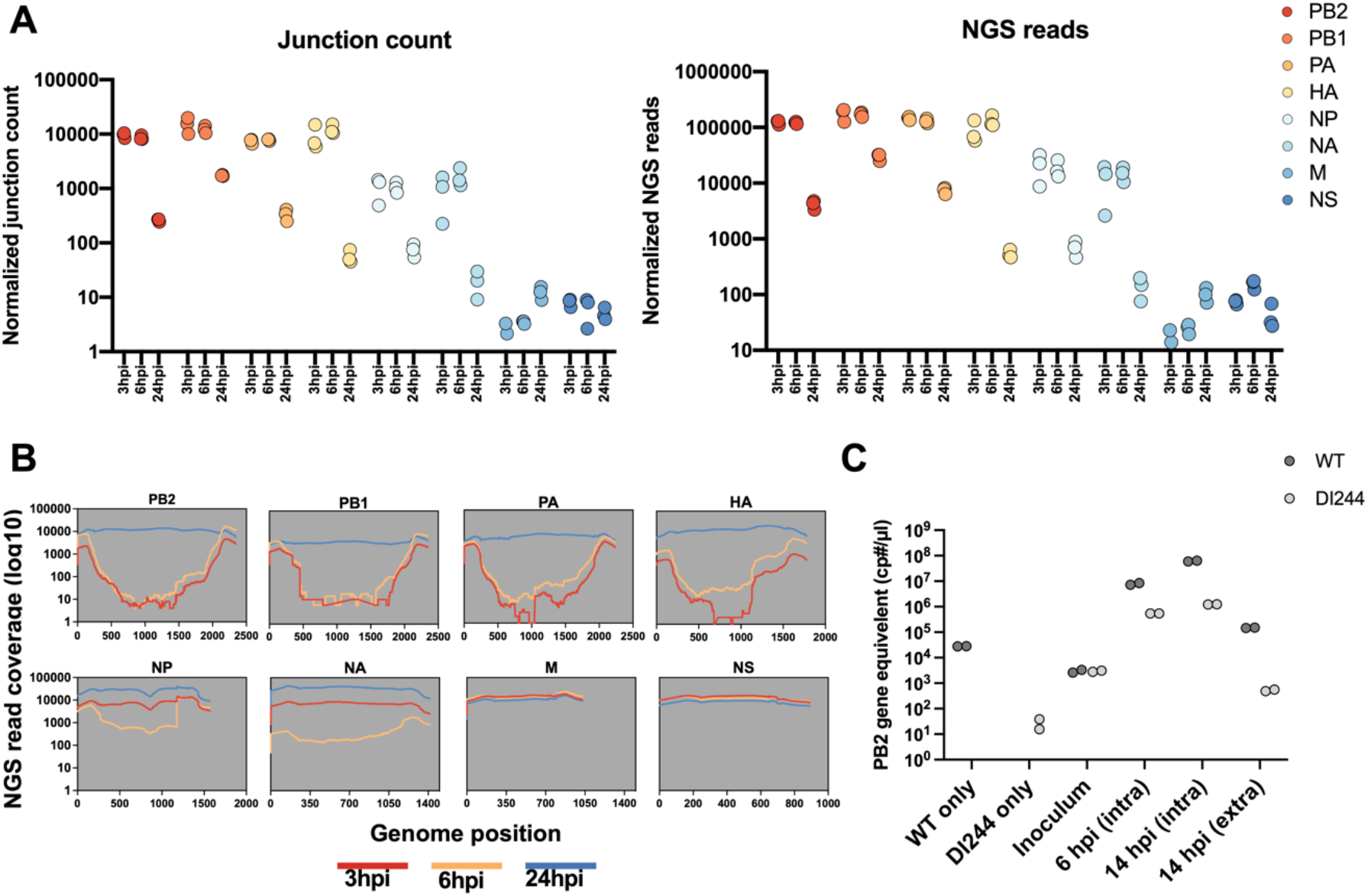
DelVGs are inefficiently packaged compared with wild type vRNAs. **(A)** Numbers of DelVG junction counts (left panel) and NGS reads (right panel) were counted for three replicates in each segment at each indicated time point, normalized to 10^6^ mapped reads. Each dot represents a replicate. **(B)** The NGS read coverage per nucleotide was plotted for each segment at each time point (the average coverage of the three replicates per position). **(C)** Competition between WT and DI244 PB2 segments during infection and packaging. MDCK-SIAT1 cells were co-infected with WT PR8 and DI244 at a 1:1 ratio to a final MOI of 20 PB2 gene equivalents/cell under single cycle conditions. Absolute copy numbers of WT and DI244 PB2 segments were quantified in by RTqPCR within either cell lysates (intracellular), supernatant (extracellular), or inoculum mixture at the indicated timepoints post-infection. PR8-only and DI244 only controls are also shown. Data represent absolute values from individual cell culture wells and are representative of two independent experiments.

While normalized DelVG junction count numbers were highly similar for individual segments between 3 hpi and 6 hpi, we observed a 10-100 fold decrease in normalized junction counts by 24 hpi for segments 1-6 **(Fig 4A, left panel)**. The same trend was also observed when we compared DelVG read support across timepoints **(Fig 4A, right panel)**. These data suggested that DelVGs constitute a much smaller proportion of viral RNAs packaged into virions, compared with what is present within the infected cell.

To confirm this observation, we examined the per-nucleotide read coverage for each segment. Since DelVGs are missing large internal regions of the genomic RNA sequence but still retain the segment termini, they produce a characteristic “devil horns” read coverage pattern, where read coverage is substantially lower in the middle of gene segment compared with the termini (25). This pattern was obvious in intracellular samples from 3 hpi and 6 hpi for segments 1-4, to a lesser extent for segments 5 and 6, and absent from segments 7 and 8, all consistent with the observed variation in abundances of DelVGs associated with each segment **(Fig 4B)**. In contrast, read coverage was consistent across the length of all eight segments in extracellular samples from 24 hpi, suggesting that DelVGs are much less abundant within this population of viral RNA.

To directly quantify the relative packaging efficiencies of DelVGs and WT vRNAs, we performed an *in vitro* competition assay between WT PR8 and a recombinant stock of DI244, a well characterized DIP/DelVG that harbors a 1946 nucleotide internal deletion in the PB2 segment (26, 27), using qPCR primer/probe-sets specific for either the WT or DI244 versions of the PB2 gene segment **(Fig 4C)**. We co-infected MDCK-SIAT1 cells with a 1:1 ratio (based on PB2 gene equivalents) of WT PR8 and DI244 at a combined MOI of 20 PB2 gene equivalents/cell under single cycle conditions where secondary spread was not permitted. We confirmed that the absolute copy numbers of both WT and DI244 PB2 gene equivalents present in the inoculum were equivalent using RTqPCR.

Quantification of both WT- and DI244-derived PB2 genome equivalents within infected cells revealed that WT PB2 was ∼50-fold more abundant than DI244 PB2 by 14 hpi, indicating that the WT PB2 segment enjoyed a significant replicative advantage over DI244. Quantification of extracellular RNA at the same timepoint (14 hpi) revealed an even more pronounced, ∼280-fold advantage for the WT PB2 segment compared with DI244 **(Fig 4C)**. Collectively, these data suggest that DelVGs make up a much smaller fraction of viral RNAs that get packaged into virions compared with what is present within the infected cell, consistent with a packaging defect relative to WT vRNAs.

### No significant differences in deletion breakpoints between intracellular and packaged DelVGs

The discrepancies in both proportional abundance and distribution amongst the segments that we observed between intracellular and extracellular DelVGs suggested the existence of a significant bottleneck limiting packaging of DelVGs relative to wild type vRNAs. We hypothesized that this might be due to the potential loss of sequences required for efficient packaging during the formation of some DelVGs. If true, we expected that specific regions of gene segments required for maximal packaging efficiency would be retained within packaged extracellular DelVGs, but largely missing from intracellular DelVGs.

To test this, we first examined the positions of deletion junctions from intracellular (6hpi) and extracellular (24 hpi) DelVGs **(Fig 5A)**. As we and others have previously reported for packaged extracellular DelVGs, deletion junctions from both populations clustered within clear hot spots near the segment termini (25, 28, 29). The extracellular DelVG junction distribution appeared narrower than that of the 6 hpi intracellular DelVGs, but that could be an artifact of the greater overall abundance of intracellular DelVGs. To more rigorously evaluate whether the locations of deletion junctions differed between intracellular and extracellular DelVGs, we used a cumulative score method that allowed us to examine the proportional distribution of deletion junctions as a function of nucleotide position **(Fig 5B)**. This approach clearly demonstrated that the distribution of deletions is statistically indistinguishable between intracellular and extracellular DelVGs (P=0.9; t-test), suggesting that DelVGs that retain specific features or sequences are not selected for during the packaging process. Finally, we asked whether the sizes of DelVGs varied between the three time points tested and again found no significant difference (p=0.57; t-test) **(Fig 5C)**, indicating that DelVG size is not a determinant of packaging efficiency.

**Fig 5.**
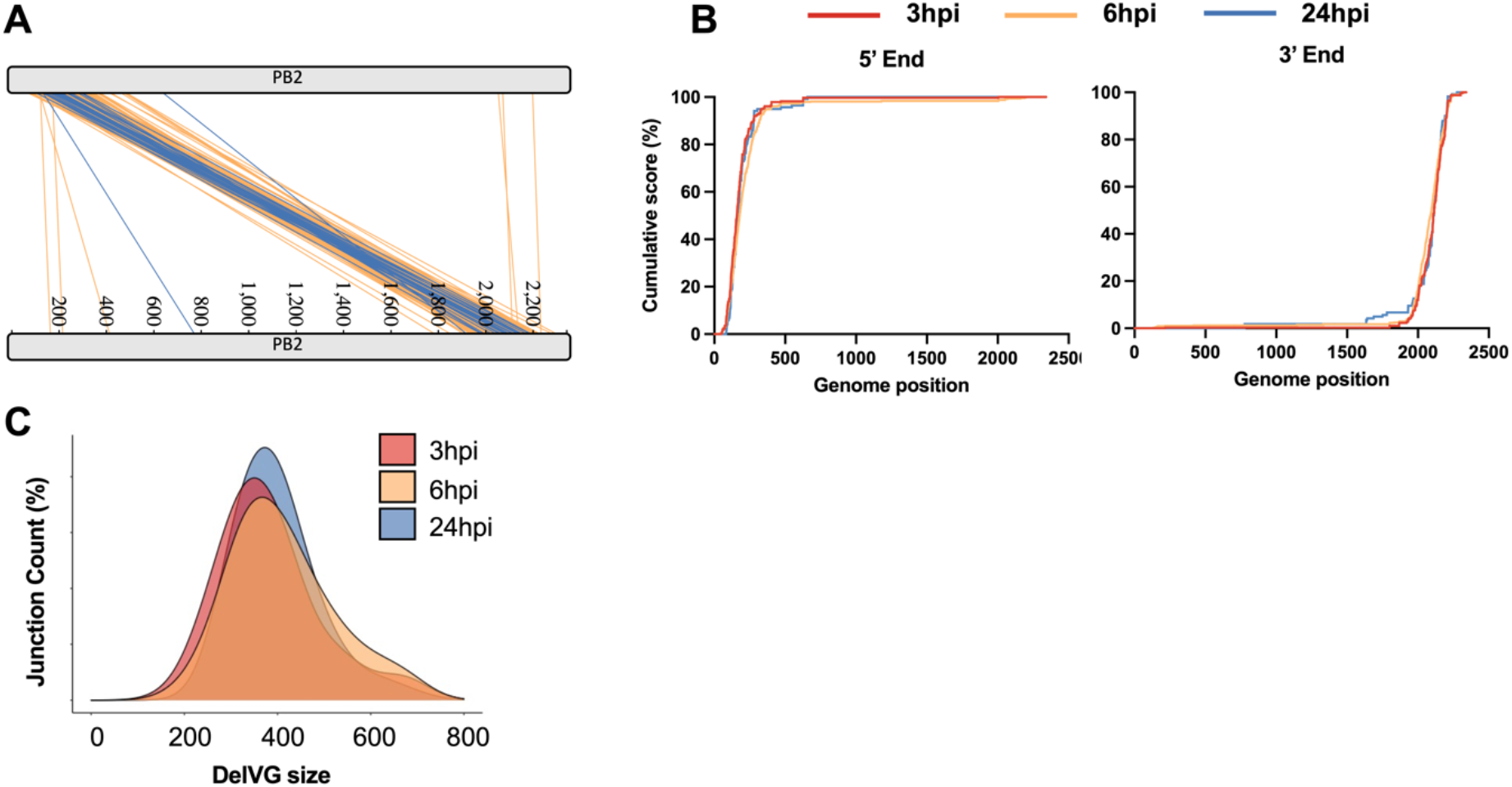
No differences in deletion size or breakpoint locations between intracellular and packaged DelVGs. **(A)** PB2 DelVG deletion junction sites at 6 and 24 hpi were mapped to the genome position with each line representing a distinct deletion and the colors indicating the time point (orange = 6 hpi, blue = 24 hpi). **(B)** Plots show the cumulative occurrence of DelVG deletions as a function of gene segment position. Cumulative score was calculated by starting at zero at the end of the segment and then adding a score of 1 at each nucleotide where a unique junction breakpoint occurred. Scores were normalized by calculating the percentage of final value reached at each position. **(C)** DelVG size distributions (PB2 segment only shown) from samples collected at the indicated time points were plotted using R density plot (ggplot package). The area under the curve is equal to 100 percent of all probabilities.

## DISCUSSION

Despite recent improvements in our fundamental understanding of the structure and function of the IAV polymerase complex, we still don’t know how or why DelVGs and DIPs form during IAV replication (11, 30–33). Here, we used a robust combined NGS/analysis pipeline to analyze the first wave of DelVGs that form within cells during infection, thus providing the first unbiased view of *de novo* DelVG production by the IAV replicase. We compared these intracellular DelVGs to the population of DelVGs that get packaged into virions, revealing a significant bottleneck in DelVG packaging relative to wild type vRNAs. Our data contradict to the dogma that DelVGs outcompete wild type vRNAs for packaging and suggest that the accumulation of DIPs within IAV populations must occur through other mechanisms.

In agreement with previous studies, we found that the majority of extracellular DelVGs junctions were derived from the three polymerase segments (24). Within intracellular DelVG populations however, the abundance of HA-derived DelVGs was comparable to that of the polymerase-derived DelVGs, suggesting that DelVGs from segments 1-4 are generated at similar rates but that the HA segment packaging efficiency is much more sensitive to deletions than the polymerase segments. It has been suggested that this bias in DelVG formation across the segments is a function of segment length, potentially because DelVGs derived from longer segments have a greater length differential compared with their wild type parents, resulting in a more pronounced replication advantage (3, 34). Our findings that the enrichment of DelVGs from segments 1-4 is already apparent by 3 hpi, suggests that this bias is emerging from the formation process rather than replication. More work is needed to identify the specific determinant(s) that influence the uneven distribution of DelVG formation across genome segments.

DelVG formation is thought to occur when the viral replicase pauses synthesis of the daughter vRNA or cRNA but continues processing along the template and re-initiates synthesis at a downstream site on the same template (3, 35). This process is not completely random, as the vast majority of deletions start and stop within hotspots near the segment termini, and individual segments vary greatly in DelVG formation (24, 25, 36). We observed the same distribution in intracellular DelVGs, indicating that these hotspots reflect that what is produced by the viral RdRp and are not biased by selection in the packaging process. Altogether, our data suggest that specific regions of the viral genome are uniquely prone to DelVG formation, for reasons that are still not understood.

It has been suggested that the polymerase translocation is promoted by the presence of a direct sequence repeat and a specific nucleotide composition at the junction site (6, 15, 24). We demonstrate significant enrichment of direct sequence repeats and A/U nucleotides at DelVG deletion junctions, along with C/G nucleotides downstream of the 5’ deletion breakpoint. Interestingly some of these nucleotides are located within the regions that are not retained in DelVG final product (R2 and R3 in Fig 3), suggesting possible roles for the sequence composition both upstream and downstream of junction sites at both ends of the viral genome. These specific template sequence features likely enhance the probability of RdRp translocation occurring, however these features are not absolutely required, as large numbers of DelVG junctions that lack flanking direct repeats or A/U bases can easily be observed. There also appears to be a significant degree of stochasticity in the specific nucleotides at which deletions form, based on the limited degree of overlap in breakpoint locations between replicates.

DelVGs/DIPs are known for their ability to inhibit the replication of WT virus, and it is widely believed that this effect is partially driven by DelVGs outcompeting WT vRNAs for packaging (37). Our data strongly suggest that the opposite is true: that DelVGs are inefficiently packaged into virions compared with WT vRNAs. This finding complicates our understanding of how DIPs outcompete WT virus at the population level, suggesting that other advantages must help DelVGs/DIPs offset their packaging deficiencies.

Given the similar characteristics of deletions between intracellular and extracellular DelVG populations, it appears that the effects of DelVG-associated deletions on packaging efficiency are independent of deletion size or location. Our data suggest that all DelVGs generated by the IAV RdRp are lacking some determinant required for maximal packaging. IAV genome packaging is governed by multiple, discontinuous regions that act in *cis* and in *trans* to facilitate the efficient and selective incorporation of a single copy of each genome segment into the vast majority of virions (29). Each segment contains packaging and bundling sequences that span both coding and non-coding regions of the segment termini (38–42). Beyond the well-described packaging signals in the segment termini, additional packaging determinants exist within the interiors of some segments (43–46). We observed no enrichment of DelVGs that retained longer stretches of the segment termini, suggesting the decreased efficiency of DelVG packaging is due to the loss of internal packaging determinants.

Our analysis of the initial wave of DelVGs produced *de novo* during IAV infection provides critical insights into the formation and packaging of both DelVGs and WT RNAs. This approach generated a detailed portrait of the full range of DelVG produced by the IAV replicase, allowing us to show that DelVGs are inefficiently packaged into particles relative to WT, contrary to dogma. Additionally, we showed that DelVG formation is not influenced by the segment length and partially influenced by sequence context. Several fundamental questions remain, however, including the specific mechanism that triggers DelVG formation and the factors that cause DelVGs to only form at defined hotspots and in higher frequency within some segments than the others.

## MATERIALS AND METHODS

### Virus and cells

MDCK-SIAT1 and HEK293T cells were grown in minimal essential medium (MEM) plus GlutaMax (Gibco), supplemented with 8.3% fetal bovine serum (Seradigm), at 37°C and 5% CO2. Recombinant A/Puerto Rico/8/1934 (PR8) was generated from HEK293T cells through standard influenza virus 8-plasmid reverse genetics transfection. Undiluted transfection supernatants were directly inoculated onto MDCK-SIAT1 cells, and supernatants were harvested at first signs of cytopathic effect (CPE) to generate seed stocks. Working stocks of virus were generated by infecting MDCK-SIAT1 cells with seed stock at an MOI of 0.0001 TCID50/cell and harvesting 48 hpi. Supernatant was clarified at 3000 RPM for 5 minutes and 500 μL aliquots were stored at -70°C.

### Generation of DelVG through high MOI infection

Confluent MDCK-SIAT1 were infected in triplicate with PR8 at an MOI of 10 TCID50/cell. To harvest intracellular viral RNA at 3 and 6 hpi, cells were washed twice with PBS, and RNA was extracted using the QIAgen RNeasy kit following the manufacturer’s instructions. To extract extracellular RNA from packaged virions, supernatant was collected from infected cells at 24 hpi. After clarification, 140µl of supernatant was used for RNA extraction using the Qiagen QIAamp viral RNA mini kit per the manufacturer’s instructions. All RNA was stored at -70°C.

### Viral cDNA amplification and sequencing

Universal RT-PCR was performed on all the samples using a previously described method (17, 47). Briefly, 3µL of RNA was mixed with 1µL (2µM) MBTUni-12 primer + 1µL (10µM) dNTPs + 8µL dH2O. Reactions were incubated as previously described, and finally 1µL SuperScript III RT (Invitrogen), 4µL of 5X First-Strand Buffer, 1µL of DTT, 1µL RNase-in (Invitrogen) were added before the final incubation. From this, 5µl was used with 2.5µL (10µM) of each universal primer, 0.5µL Phusion polymerase (NEB), 10µL - 5x HF buffer, 1µL (10mM dNTPs mix), and 28.5µL dH2O. The PCR conditions used were 98°C (30 s) followed by 25 cycles of 98°C (10 s), 57°C (30 s), and 72°C (90 s); a terminal extension of 72°C (5 min); and a final 10°C hold. Finally, PCR amplicons were purified using the Qiagen QIAquick PCR Purification Kit and sequenced with paired-ends 2×250nt reads on an Illumina MiSeq using V2 chemistry (17).

### Generation of recombinant DI244

We synthesized (Integrated DNA technologies, Inc.) and cloned the full-length DI244 sequence (NCBI # L41510.1) into the pDZ vector and transfected it along with 7 plasmids encoding WT versions of segments 2-8 plasmids into PB2-expressing HEK293 cells (HEK293-PB2), using the standard 8-plasmid reverse genetics approach. Briefly, 60 to 90% confluent PB2-expressing HEK293 cells were transfected with 500 ng of various plasmids (pDZ::PB2-DI244, pDZ::PB1, pDZ::PA, pDZ::HA, pDZ::NP, pDZ::NA, pDZ::M, and pDZ::NS) by using JetPrime (Polyplus) according to the manufacturer’s instructions. Under these conditions, no WT virus is generated due to the absence of WT PB2 segment. Transfection supernatant was used to infect PB2-expressing MDCK cells (MDCK-PB2) for 48 hrs to generate a seed stock. HEK293-PB2 and MDCK-PB2 cells were kindly provided by Dr. Stefan Pöhlmann and are described in this reference (5). DI244 seed stock was titered on MDCK-PB2 cells and used to infect MDCK-PB2 cells at an MOI of 0.001 TCID50/cell for 72 hrs to generate a working stock. Deep sequencing of the DI244 working stock confirmed the presence of the expected DI244 deletion junction and the absence of WT PB2 segment (data not shown).

### RTqPCR quantification of DI244 and WT PB2 gene segments

We designed and optimized specific primer/probesets (**Table S1**) specific for either DI244 or WT PB2 using the IDT PrimerQuest webtool and validated efficiency and specificity using serial dilutions of plasmids encoding either WT PB2 or DI244. Viral RNA was extracted from cells or virions as described above and used to synthesize cDNA using the universal primer described above and *Verso* cDNA synthesis Kit (Thermo Fisher). 3µl of RNA was mixed with 8µl H20, 4µl 5X cDNA Synthesis Buffer, 2µl dNTP Mix (5 mM each), 1µl universal primer (10µM), 1µl RT Enhancer, 1µl Verso Enzyme Mix, before incubating for 50 minutes at 45°C. After this, 1µl of the of cDNA product was mixed with 7µl H20, 1µl forward primers (18µM), 1µl reverse primer (18µM), 1µl specific probe (5µM), and 10µl TaqMan Fast Advanced Master Mix (Thermo Fisher). The qPCR conditions used were: 50°C (2min), 95°(2 min), followed by 40 cycles of 95°C (1 s) and 61°C (20 s) using a qPCR QuantStudio 3 thermocycler.

**Table S1:**
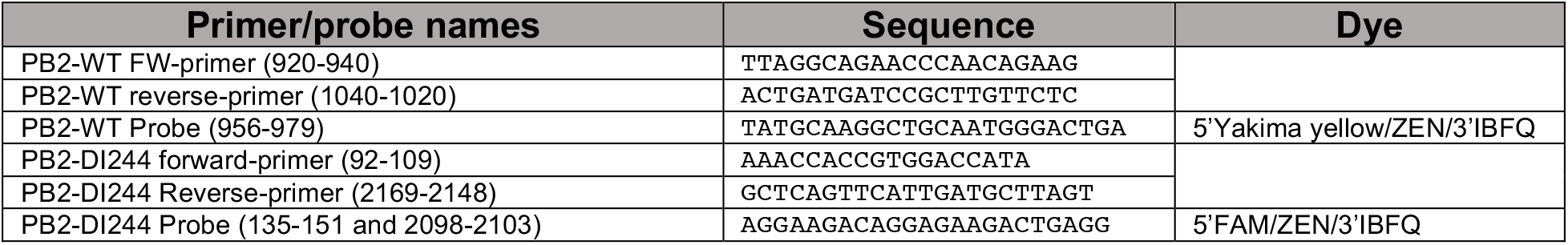
qPCR primers and probes used for the competition assay

### *In vitro* competition assay

MDCK-SIAT1 cells were coinfected in duplicate at an MOI of 10 PB2 gene equivalents/cell with a 1:1 ratio of WT PR8 and DI244 (ratio based on PB2 gene equivalents). Fractions of the inoculum mixture both before (0 hpi) and after the 1 hour adsorption phase were set aside for RTqPCR. After adsorption for 1 hour at 4°C, inoculum was removed, cells were washed, and MEM+FBS was added to cells. At 3 hpi, neutralizing anti-HA monoclonal antibody H36-26 was added to each well to a final concentration of 25ug/mL to block secondary spread. Cellular RNA was extracted from duplicate wells of cells at 6 hpi and at 14hpi using the QIAgen RNeasy kit following the manufacturer’s instructions. At the 14hpi timepoint, viral RNA was also collected from supernatants using the Qiagen QIAamp viral RNA mini kit per the manufacturer’s instructions. Control infections with only WT PR8 or only DI244 were also performed in parallel.

### Sequencing analysis of deletion junctions

Raw sequencing reads were fed into our DI-detection pipeline for junction detection and characterization (17). Briefly, the pipeline uses set of algorithms including Bowtie (48) and ViReMa (49) to perform two major steps. First, filtering the NGS reads based on quality and origin (i.e. cellular or viral). Second, mapping and detecting the deletion junction and reporting the start and end sites for each junction in each gene segment, along with the number of NGS reads detected for each junction. The pipeline is accessible at: https://github.com/BROOKELAB/Influenza-virus-DI-identification-pipeline. NCBI accession numbers for the PR8 reference genome used in this study: PB2=AF389115.1, PB1=AF389116.1, PA=AF389117.1, HA=AF389118.1, NP=AF389119.1, NA=AF389120.1, M=AF389121.1, NS= AF389122.1

To account for sequencing read coverage variations between libraries, we normalized the junction count and their NGS reads to 10^6^ mapped reads per library per segment. For the junction count, the number of junctions per segment was multiplied by 10^6^ and divided by the number of NGS reads aligned to the WT gene of the given segment in a given library. The same was done to normalize for the NGS reads, where the junctions’ read number per segment was multiplied by 10^6^ and divided by the number of NGS reads aligned to the WT genome of the given segment plus the number of reads mapped to the junctions.

The direct repeat sequence lengths were extracted from the output file “Virus_Recombination_Results.txt” generated by ViReMa algorithm. For the random control, the junction sites were randomized using Excel function “=RANDBETWEEN()” based on the actual sequence range and number detected in the 6 hpi population. Next, a custom Perl code was used to extract their sequences from the corresponding PR8 gene segment (PB2 and HA). Finally, the direct repeat lengths were extracted and compared to the real samples.

To analyze the nucleotide composition at the junction site, we analyzed the sequences flanking the junction sites for enrichment of specific nucleotides. There are four possible sequence regions that possibly involved in promoting the polymerase translocation, two regions flanking the junction from each site, we numbered them region 1-4 **(Fig 3)**. From these regions only, R 1& 4 are retained within the DelVG final product, while R 2 & 3 are not, and their potential importance stems from their physical proximity to the junction. A custom Perl code was used to extract 4 nucleotides plus the junction from each region from all the detected DelVG in the three replicates of the 6 hpi population. Next the WebLogo platform (50) was used to measure the percentage occurrences of each nucleotide at each position. To decide whether the percentage occurrence differed from what would be expected in the absence of nucleotide enrichment, we generated a random control samples the same way as in the direct repeats. Finally, the two samples: observed/experimental and expected/computational were compared using Anova Two-way. To confirm the validity of this approach we found the same results when repeated the analysis for 3 hpi time point and different segments. Additionally, we found no significant difference at all nucleotide position upon comparing two random samples, three replicates each (data not shown).

The percentage length of each segment was calculated based on the total genome length 13585nts (*e*.*g*. PB2=17.2% and NS=6.5%). Next, the percentage length of each segment was used to calculate the number of junctions per segment based on the total normalized number of junctions of each sample (expected). Finally, these values were compared to the observed values from the actual experiments.

### Data availability

All NGS data sets generated in this study can be found under BioProject accession number `PRJNA725907`.

## Acknowledgment

We are grateful to other members of the lab for helpful comments and critical readings of the manuscript, as well as to Dr. Tanja Laske from Max Planck Institute and Dr. Prerna Arora from Georg-August-University Göttingen for helpful discussion. This work was generously funded by the Defense Advanced Research Projects Agency under contract DARPA-16-35-INTERCEPT-FP-018. A.t.V. is supported by a joint Wellcome Trust and Royal Society grant 206579/Z/17/Z.

